# Factors underpinning discrepant findings from reviews on the same topic: A systematic analysis of Cochrane and non-Cochrane reviews

**DOI:** 10.1101/574046

**Authors:** Claudia Hacke, David Nunan

**Author notes:** **Corresponding Author:** Dr. Claudia Hacke, University Medical Center Schleswig Holstein, Campus Kiel, Arnold-Heller-Straße 3, 24105 Kiel, Germany. ORCID: 0000-0002-3941-5129.

## Abstract

**Objective:** To explore factors underpinning discrepancies in reported pooled effect estimates from Cochrane and non-Cochrane systematic reviews answering the same question.

**Study Design and Setting:** We observed discrepant pooled effects in 23 out of 24 pairs of meta-analyses from Cochrane and non-Cochrane systematic reviews answering the same question. Here we present the results of a systematic assessment of methodological quality and factors that explain the observed quantitative discrepancies. Methodological quality of each review was assessed using AMSTAR (Assessing the Methodological Quality of Systematic Reviews). Matched pairs were contrasted at the macro- (review methodology), meso- (application of methodology) and micro- (data extraction) level and reasons for differences were derived.

**Results:** All Cochrane reviews had high methodological quality (AMSTAR 8-11), whereas the majority (87.5%) of non-Cochrane reviews were classified as moderate (AMSTAR 4-7). Only one pair included exactly the same studies for their respective meta-analyses but there was still a discrepancy in the pooled estimate due to differences in data extraction. One pair did not include any study of its match and for one pair the same effect estimates were reported despite inclusion of different studies. The remaining pairs included at least one study in their match. Due to insufficient reporting (predominantly affecting non-Cochrane reviews) we were only able to completely ascertain the reasons for discrepancies in all included studies for 9/24 (37.5%) pairs. Across all pairs, differences in pre-defined methods (macro-level) including search strategy, eligibility criteria and performance of dual screening could possibly explain mismatches in included studies. Study selection procedures (meso-level) including disagreements in the interpretation of pre-defined eligibility criteria (14 matches) were identified as reasons underpinning discrepant review findings. Comparison of data extraction from primary studies (micro-level) was not possible in 13/24 pairs as a result of the non-Cochrane review providing insufficient details of the studies included in their meta-analyses. Two out of 24 pairs completely agreed on the numerical data presented for the same studies in their respective meta-analysis. Both review types provided sufficient information to check the accuracy of data extraction for 8 pairs (45 studies) where there were discrepancies. An assessment of 50% (22 studies) of these showed that reasons for differences in extracted data could be identified in 15 studies. We found examples for both types of review where data presented were discrepant from that given in the source study without a plausible explanation.

**Conclusion:** Methodological and author judgements and performance are key aspects underpinning poor overlap of included studies and discrepancies in reported pooled effect estimates between topic-matched reviews. Though caution must be taken when extrapolating, our findings raise the question as to what extent the entire meta-analysis evidence-base accurately reflects the available primary research both in terms of volume and data. Reinforcing awareness of the application of guidelines for systematic reviews and meta-analyses may help mitigate some of the key issues identified in our analysis.

**What is new?:** Key findings

- Non-Cochrane reviews were of a lower overall methodological quality compared with Cochrane reviews. Discrepant results of meta-analyses on the same topic can be attributed to differences in included studies based on review author decision, judgements and performance at different stages of the review process.

**What this adds to what was known?:**

- This study provides the most robust analysis to date of the potential methodological factors underpinning discrepant review findings between matched meta-analyses answering the same question. Assessing differences between reviews at the macro-, meso-, and micro-levels is a useful method to identify reasons for discrepant meta-analyses at key stages of the review process.

**What is the implication and what should change now?:**

- There is a need for a standardised approach to performing matched-pair analysis of meta-analyses and systematic reviews answering the same question. Our paper provides a base for this that can be refined by replication and expert consensus.

## 1. Introduction

There has been an exponential growth in published systematic reviews in recognition of their pivotal role as part of evidence-based practice and guideline development [1]. With that, however, comes an increasing number of reviews on similar topics and potential for overlap and duplication [2] potentially contributing to research waste [3]. The need to differentiate well from poorly conducted reviews will become increasingly important for users of systematic reviews including evidence-based practitioners, policy makers and guideline developers.

There is empirical evidence indicating that Cochrane reviews and systematic reviews published outside the Cochrane Collaboration differ considerably in their methodologically quality, and can provide discrepant results and conclusions, even if answering the same question over a similar time frame [4–10]. Accordingly, in our matched-pair analysis of Cochrane and non-Cochrane reviews we identified, for the first time, inconsistencies in pooled effect estimates between meta-analyses examining the role of physical activity interventions for prevention and treatment of major chronic diseases (Hacke & Nunan, 2019). These results are consistent with those of a recent analysis of pharmacological interventions which showed generally poor overlap of included studies between matched pairs of Cochrane and non-Cochrane reviews [9].

The evidence to date, however, has not adequately addressed the potential methodological factors underpinning observed discrepancies among topic-matched reviews. Useem et al. [9] suggested different search strategies and inclusion and exclusion criteria as potential explanations for discrepancies but did not explicit evaluate these or other factors, instead recommending them for future work.

To complete the quantitative analysis of matched Cochrane and non-Cochrane paired reviews of physical activity interventions in our sister paper (Hacke & Nunan, 2019), here we aimed to qualitatively explore the methodological factors underpinning the discrepancies we observed. Insights gained from this study may help to understand and evaluate discrepant findings from systematic reviews and meta-analyses and reinforce the need for adherence to sound methodological practice and reporting.

## 2. Material and Methods

Our quantitative study found discrepancies in reported pooled effect estimates for 24 pairs of meta-analyses, each obtained from a Cochrane review and non-Cochrane review matched according to the same intervention, condition and outcome, and published within five years of each other (Hacke & Nunan, 2019). In order to explain the observed discrepancies we firstly evaluated the methodological quality using the AMSTAR (Assessing the Methodological Quality of Systematic Reviews) tool [11]. Accordingly, we ranked reviews as high quality (AMSTAR score 8-11), medium quality (scoring 4-7), or of low quality (scoring 0-3) [11, 12].

We then evaluated potential factors underpinning the observed quantitative discrepancies. This was done firstly at the macro-level by assessing the specific methodologies employed in both review types. This included:

- A-priori defined criteria: protocol mentioned and accessible
- Details of search: Number of databases, years of coverage, search period, Key words/MESH terms, full search strategy, supplementary search strategies
- Eligibility criteria: Study design, publication status, language
- Data abstraction: Double screening and extraction, data synthesis
- Results: List of included and excluded studies

We then assessed discrepancies at the meso-level including overlap of primary studies, reasons reported for study exclusion, with the use of lists of excluded studies if available and considering the search period of each review of a matched pair. Where no list of excluded studies was available or a study was not listed as excluded, and no other reason was apparent that would justify the exclusion of a specific study from the meta-analysis, we stated the primary study as “not found”, meaning that there is no evidence that this study was included in the list of relevant studies in the review screening process. Subsequently, reasons for discrepant findings of a matched pair were classified based on discrepancies that could be attributed to different pre-defined inclusion criteria, or factors that were related to discrepancies in the interpretation of pre-defined inclusion criteria, the decision on eligibility of study characteristics and other factors (see results “Comparison of reviews at the meso- and micro-levels”). Finally, we explored reasons for discrepancies at the micro-level between matched-pair pooled effect estimates. This included, where available, the discrepancies in presented data (number of participants, means and SD,) of each primary study that was found and included in matched-pair meta-analyses. We checked ^~^50% of the studies with discrepant data presented.

## 3. Results

### 3.1 Quality of reviews

The methodological quality of included reviews is presented in Figure 1. The overall AMSTAR score was 9.5 (IQR 9-10) for the 24 Cochrane reviews and 5 (IQR 4-6) for the non-Cochrane reviews. All Cochrane reviews were judged high quality (≥ 8) according to AMSTAR assessments. In contrast, only one non-Cochrane review were judged high quality, two were of low quality (AMSTAR 0-3) and 21 were classified as moderate quality (AMSTAR 4-7). Stating a potential for a conflict of interest (91.7%) and the consideration of publication bias (37.5%) were most likely to be missing in Cochrane reviews, but were more often missed in non-Cochrane reviews (95.8% and 50.0% respectively). Non-Cochrane reviews were highly likely to not refer to a priori published research objectives (91.7%) and to exclude studies based on publication type or language (91.7%). The majority of both review types provided lists and describes the characteristics of included studies, however, in some pairs (7/24, 29.2%) the non-Cochrane review listed the number of studies for a given intervention or outcome, but did not specify which of those articles pertained to a specific meta-analysis. Similarly, only one-quarter of non-Cochrane reviews (6/24, 25%) but all Cochrane reviews (24/24, 100%) provided a list of excluded studies with reasons.

**Fig.1.**
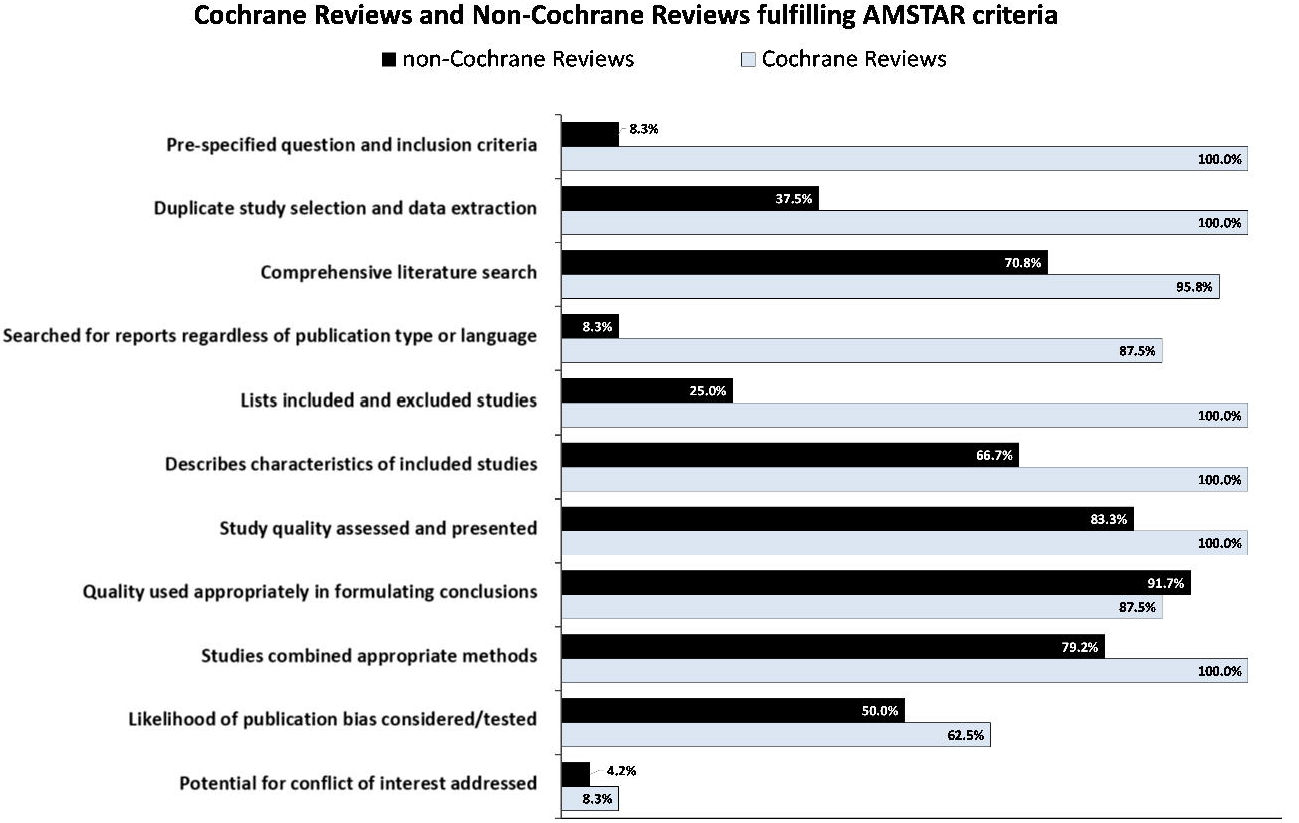
AMSTAR assessments for included Cochrane and non-Cochrane reviews (n=24 pairs). Data presented are the percentage of reviews fulfilling each criterion.

### 3.2 Comparison of review characteristics at the macro-level

All Cochrane reviews reported a full electronic search strategy in their methods compared to just under half (11/24, 45.8%) for non-Cochrane reviews (see Appendix 1). Cochrane reviews reported searching more databases than did non-Cochrane reviews, a median of 7.5 (IQR 5.25-10.0) electronic databases compared with 6.5 (4.25-11.0) for non-Cochrane reviews. All Cochrane reviews and all but one non-Cochrane review reported on the years covered for their database search, with Cochrane reviews reporting a search period 9 months longer on average than their matched non-Cochrane review. The majority of reviews reported that the search was complemented by at least one supplementary search strategy (45/48, 93.8%), with searching for grey literature in Cochrane reviews (24/24, 100%) and checking for reference lists in non-Cochrane reviews (18/24, 75%) as the most often applied supplementary strategies. However, Cochrane reviews were more likely to supplement their search with additional sources. Only one Cochrane review (1/24, 4.2%) but half of the non-Cochrane reviews (12/24, 50.0%) excluded studies based on language. Inclusion of both published and unpublished studies was more likely in Cochrane (23/24, 95.8%) compared with non-Cochrane reviews (11/24, 45.8%).

In only about one-third of the non-Cochrane reviews (9/24, 37.5%) was it stated that both selection of studies and extraction of data were performed by two (or more) independent authors, while all Cochrane reviews performed dual screening and extraction.

Overall, a random-effects model was applied in just over half of the meta-analyses performed (28/48, 58.3%), with differences in models applied between Cochrane and non-Cochrane found in 6/24 (25%) of matched-pairs. The majority of both review types provided lists of included studies, however, in some pairs (6/24, 25%) the non-Cochrane review listed the number of studies for a given intervention or general outcome, but did not specify which of those articles pertained to a specific meta-analysis. Similarly, only one-quarter of non-Cochrane reviews (6/24, 25%) but all Cochrane reviews (24/24, 100%) provided a list of excluded studies with reasons.

### 3.3 Comparison of reviews at the meso- and micro-levels

We contrasted each Cochrane and non-Cochrane review of a matched pair to examine the overlap of primary studies. Reasons for study exclusion were assessed taking into consideration the review list of excluded studies and the period of literature search.

With regard to the assessment on the meso-level, only one pair included exactly the same studies but did not report the same pooled effect estimate. Overall, in one-third of matches (9/24, 37.5%) we were able to find all of the reasons for differences in study overlap. However, in the other two-thirds of the matched pairs (15/24, 62.5%) one or more studies (maximum = 25 studies) that were included in one review could not be found in its match for reasons unexplained. Plausible explanations might be that a review did not retrieve a study in its original search or a study was retrieved but discarded at the title and abstract or the full-text eligibility screening stage without information being reported. This latter issue was less likely to be relevant to Cochrane reviews due to their mandatory reporting of excluded studies (at full-text screening stage) and supporting reasons. As can be seen from the table in Appendix 2, half of the Cochrane reviews (12/24, 50%) included studies that were out of the search period of their non-Cochrane match. By contrast, only 5 non-Cochrane reviews included studies that could not be found by their Cochrane match due to a shorter search period.

#### 3.3.1 Differences in study overlap based on pre-defined criteria

About one-third of the matched pairs (14/24, 58.3%) were found to disagree in their included studies due to discrepancies in their pre-defined selection criteria with regard to study design (e. g. RCTs with or without quasi randomisation), condition (e. g. wider inclusion criteria of participants with risk factors for diabetes), intervention characteristics (e. g. group format ratio, follow-up period, drop-out rate), control intervention (e. g. usual care without vs. with or without exercise advice; no intervention vs. no intervention and intervention of different dose) (see Appendix 2).

#### 3.3.2 Differences in study overlap based on other factors related to subjective decisions

The majority of reviews screened and found most of the studies of its match but did not include them in their meta-analysis (see Appendix 2). Explanations could be categorized into those where Cochrane and non-Cochrane reviews disagreed in the interpretation of pre-defined inclusion criteria and other factors (see Appendix 3 for a detailed list of reasons with examples).

Interpretation of pre-defined inclusion criteria including:

- Study design: both reviews of a matched pair reported inclusion of RCTs, but one of them considered that a study was not an RCT when the other considered it was.
- Intervention: both reviews of a matched pair reported inclusion of studies of any exercise interventions, but one of them considered a study as providing an exercise intervention when the other did not, or that co-interventions were provided to the control and the other considered they were not; or that a specific type of exercise (e.g. interval training) was performed as described but the other considered it was not; or that one review considered that the control group received an additional active intervention when the other did not.
- Control: both reviews of a matched pair reported inclusion of non-exercise controls only, but one of them considered that the control group received exercise where the other did not; or that the control intervention differed to the standard recommendation but the other review judged it did not differ.
- Condition: both reviews of a matched pair reported inclusion of participants with the disease of interest and considered the same study, but one reported that participants did not have the disease and the other considered they did; or both reviews reported inclusion of studies with the same minimum distribution of patients with the disease of interest, but one review judged to exclude a study as the proportion of people with the disease was too low while the other match judged to include it; or both reviews focus on the effect of an intervention and considered the same study, but one review judged to exclude it as the proportion of control patients who have attended were too low while the other match judged to include it.
- Outcome: both reviews of a matched pair reported inclusion of studies with the outcome of interest, but one review judged that a study did not measure the outcome and the other considered it did.

Other factors including:

- Availability of study data: both reviews excluded studies due to missing values/unreported data, but such data were available for a study and this was reported in one review but not the other.
- Secondary analysis of studies: both reviews considered a study as a secondary analysis but one review judged to exclude it and the other did not.
- Pooled data for several measures of outcome: both reviews focus on the same outcome but one review performed separate meta-analyses according to different measures whereas the other included all measures in one meta-analysis.
- Discrepancies in sample size and/or citations: both reviews pooled data for mortality up to 12 months, but one review gives sample size and references for overall-mortality only whereas the other review gives sample size and references for mortality up to 12 months. Or both reviews included the same outcome measure, but one review gave only sample size without stating which studies were used.

Comparisons at the micro level were not possible in 13/24 pairs as a result of insufficient reporting of data from the non-Cochrane review. One pair did not include any of the study of its match. Only 2/24 pairs completely agreed on the numerical data presented for the same studies when included in their respective meta-analysis. In the remaining 8 pairs, 45 studies included in both the Cochrane and non-Cochrane review had differences in extracted data that could be assessed. We checked ^~^50% for detailed information (22 studies). Of these 22 studies, reasons for differences in extracted data could be identified in 15 studies. In the remaining 9 studies extracted data differed for reasons unexplained, including, but not limited to, a lack of reporting by the non-Cochrane review (3 studies) or the primary study (2 studies).

We identified the following reasons for differences in extracted data for 15 studies:

- Different methodologies used to assess the effects of intervention: the primary study reported data from a 3-arm intervention study and the Cochrane Review pooled the exercise and walking group and compared with control, while the non-Cochrane used the exercise group for comparison only.
- Different outcome of interest extracted: the primary study reported the outcome of interest providing different scales and the Cochrane review reported scale 1, while the non-Cochrane review reported scale 2; the primary study reported the outcome of interest at different time points and the Cochrane review extracted data at 3 months of follow up, while the non-Cochrane review extracted data at 6 months of follow up; the Cochrane review reported posttest means, while the non-Cochrane review reported change scores.
- Different sources to obtain data: both reviews reported data from a study that was published in different sources and the Cochrane review may have used data from a doctoral thesis of 2002, while the non-Cochrane review used data from a journal publication of 2005.
- Issue of type error
- Issue of rounding

## 4. Discussion

### 4.1 Principle findings

We found that non-Cochrane reviews were of a lower overall methodological quality compared with Cochrane reviews. We also identified discrepant results of meta-analyses on the same topic can be attributed to differences in included studies based on review author judgements at different stages of the review process including search strategy, the application of inclusion and eligibility criteria, and the performance of dual screening and data extraction. Discrepant findings from meta-analyses are also attributed to differences in data abstraction and their statistical analyses. However, for a large proportion of matched reviews where we identified a difference in included studies we were unable to ascertain the reason for such differences. Though caution must be taken when extrapolating our findings to meta-analyses of differing scope, they raise the question as to what extent the entire meta-analysis evidence-base accurately reflects the available primary research both in terms of volume and data. Reinforcing awareness of the application of guidelines for systematic reviews and meta-analyses may help mitigate some of the key issues identified in our analysis.

### 4.2 Comparison with existing literature

Previous studies that have been carried out on the comparison of Cochrane and non-Cochrane reviews have been restricted to an overall comparison of methodological characteristics on the macro level [4–6, 8, 10]. Accordingly, with the use of AMSTAR tool, our findings match those observed in earlier studies demonstrating a heterogeneous methodological quality between both review types [4, 5, 7, 8, 10], with particular concerns to the use of protocols and listing included and excludes studies within non-Cochrane reviews. Important gaps between Cochrane and non-Cochrane reviews were also identified in the methods employed for literature search and study selection.

These findings support earlier assumptions by Useem et al. [9], who discussed different search strategies and/or inclusion/exclusion of studies as potential explanations for discrepancies. In our study, only 50% of non-Cochrane reviews reported searching and inclusion of studies irrespective of language and only 46% irrespective of status of publication. Limitations in these eligibility criteria point to a lack of a comprehensive literature search and are, therefore, likely related to the poor overlap of studies between Cochrane and non-Cochrane reviews and subsequently resulting in the discrepant effect estimates we observed (Hacke & Nunan, 2019).

Previous studies similar to ours have note adequately addressed the reasons for discrepancies among a topic-matched pair [9, 10]. In the present study, non-Cochrane reviews were less likely to find the included studies of its Cochrane review match. This finding may have been biased by the lack of reporting of excluded studies in non-Cochrane reviews (18/24, 75.0%), corroborating the findings of Moher [4] and limits our ability to understand which studies were given consideration as well as to judge the quality of search and justification of study selection.

Though similar in year of publication, our comparison at the meso-level confirms that one reason for discordancy was the database search period, with Cochrane reviews applying a nine-month longer search period on average. Secondly, we could anticipate discrepant results between some of the matched pairs as some already differed in their pre-defined inclusion and exclusion criteria (see table 2). It is, however, important to note that potential discrepancies resulting from these different eligibility criteria could be attributed to a more broad literature search as inclusion criteria of reviews within a matched pair were not always entirely contradicting.

Another important finding that could be derived from meso-level evaluations was the difference in subjective judgements and decisions applied by review authors. The majority of reviews found the same studies of its match, however, authors made different judgements as to their inclusion for analysis based on interpretations of selection criteria with regard to study design, the intervention, the control, the treating condition or the outcome. This led to different decisions as to which studies were considered eligible or not. In this regard, some reviews stated reasons for exclusion of studies to suggest that its matched review did not only differ in the definition but even did not adhere to its own pre-defined inclusion criteria. For example, both reviews may have stated they would include studies that employed a non-exercise control intervention only, but one of them included a study with an exercise control. The issue of inconsistencies within matched paired meta-analyses due to conflicting interpretations of eligibility criteria is an intriguing one which could be explored in further research, including surveying of or qualitative interviews with systematic review authors as well as journal editors and peer-reviewers.

With regard to macro-level comparisons, only 8.3% of matched pairs (2/24 pairs) completely agreed on the numerical data extracted from the same studies when included in their respective meta-analysis. One issue that can be observed is that that almost two-thirds of non-Cochrane reviews (64%) did not follow best practice by performing study selection and data extraction independently by at least two researchers. However, we found examples for both Cochrane and non-Cochrane reviews where it was not possible to determine how review authors arrived at the data they presented from examining the primary study.

Observed differences in methodological and reporting quality might reflect differing or nonexistent guidelines between reviews inside and outside the Cochrane Collaboration. On the contrary, with the development of the QUOROM/PRISMA Statements [13, 14] concise guidelines for systematic reviews and meta-analyses of evaluations of health care interventions are available since 1999.

In terms of the impact on the findings from the two reviews, discrepancies in methodological quality, subjective judgements and data extraction resulted in 25% of match-pair pooled effect sizes differing in their statistical or clinical interpretation the poor study overlap. This may be the result of both Cochrane and non-Cochrane reviews including a similar number of studies and participants thus ensuring similar variance around reported effect estimates. Equally it may also reflect the true effect of the intervention across different studies. Similar findings have been observed in match-pair analyses of pharmacological interventions for patients with cardiovascular disease [9].

### 4.3 Strengths and limitations of this study

Our study provides the most robust analysis to date of the methodological quality of matched Cochrane and non-Cochrane reviews and offers, for the first time, possible reasons underpinning our observation that meta-analyses from Cochrane and non-Cochrane reviews assessing the same questions show discrepancies in reported treatment effect estimates and their statistical and clinical interpretation (Hacke & Nunan, 2019). A strength of our study is the exploration of methodological factors that could explain observed differences within matched pairs that have not been addressed sufficiently in previous studies [9, 10]. Another strength is the comparative evaluation of each matched Cochrane and non-Cochrane meta-analysis on the meso- and micro level to provide a series of definite reasons that impact on the extent of study overlap and results of reviews.

A limitation of this study is, as is the case with any appraisal of methodological quality, that our judgements are based on subjective decisions. Provision of our justifications for judgements will reduce misinterpretation and enable useful debates. A further limitation was restriction of some of elements of our analyses to a small number of matched-pairs due to incomplete reporting in included non-Cochrane systematic reviews. Our assessment was of 24 matched meta-analyses, a likely small proportion of the total available. Conducting a similar analysis to that performed here on already published matched-pair studies would be a relatively simple first step towards validating our findings.

## 5. Conclusion

A-priori differences in methodology and issues relating to their application as well as data extraction were key reasons for poor overlap of included studies and differences in pooled effect estimates between matched meta-analyses answering the same question. Ascertaining reasons for differences between reviews was hampered by the lack of reporting of relevant information, particularly but not exclusively from non-Cochrane reviews. Though caution must be taken when extrapolating to different fields, our findings raise the question as to what extent the entire meta-analysis evidence-base accurately reflects the available primary research both in terms of volume and data. To improve validity and replicability of meta-analyses and systematic review findings, authors should ensure provision of justifications for judgements made and actions performed throughout the review process. Journal editors and peer-reviewers should ensure adherence to reporting guidelines as a pre-requisite to publishing.

## Data Availability

All data generated or analysed during this study are included in this published article [and its supplementary information files].

## Author contributions

**C.H.; D.N.:** Conceptualization, Methodology, Validation. **C.H.:** Formal Analysis, Data curation, Investigation, Writing-Original draft preparation, Visualization. **D.H.**: Supervision. **C.H.; D.N.:** Writing- Reviewing and Editing.

## Acknowledgements

The authors would like to thank Nia W. Roberts for her role in the original search of Cochrane Systematic Reviews.

## Supplementary Material

**Appendix 1.** Macro-level analysis.

**Appendix 2.** Meso-level analysis.

**Appendix 3.** Macro-level analysis (detailed).

